# Effects of episodic future thinking on temporal discounting: a re-analysis of six data sets using hierarchical Bayesian parameter estimation and compilation of effect sizes

**DOI:** 10.1101/2020.03.24.005892

**Authors:** Jan Peters, Stefanie Brassen, Uli Bromberg, Christian Büchel, Laura Sasse, Antonius Wiehler

## Abstract

Temporal discounting refers to the tendency of humans and many animals to devalue rewards as a function of time. Steep discounting of value over time is associated with a range of psychiatric disorders, including substance use disorders and behavioral addictions, and therefore of potentially high clinical relevance. One cognitive factor that has repeatedly been shown to reduce temporal discounting in humans is episodic future thinking, the process of vividly imagining future outcomes, which has been linked to hippocampal mechanisms in a number of studies. However, the analytical approaches used to quantify the behavioral effects have varied between studies, which complicates a direct comparison of the obtained effect sizes. Here we re-analyzed temporal discounting data from previously published functional magnetic resonance imaging (fMRI) and behavioral studies (six data sets from five papers, n=204 participants in total) using an identical model structure and hierarchical Bayesian parameter estimation procedure. Analyses confirmed that engagement in episodic future thinking leads to robust and and consistent reductions in temporal discounting with on average medium effect sizes. In contrast, effects on choice consistency (decision noise) where small and with inconsistent directionality. We provide standardized and unstandardized effect size estimates for each data set and discuss clinical implications as well as issues of hierarchical Bayesian parameter estimation.

## Introduction

Temporal discounting refers to the tendency of humans and many animals to de-value rewards as a function of the time to their delivery^1,2^. While earlier studies have focused on steep reward discounting in substance-use-disorders^3^ and behavioral addictions such as gambling disorder^4^, both steep and shallow discounting have been associated with various psychiatric and neurological disorders^5^.

In the light of these associations between temporal discounting and mental disorders, cognitive factors and interventions with the potential to attenuate discounting are of considerable clinical interest^6,7^. One such mechanism that has gained substantial empirical support in recent years is episodic future thinking, that is, the ability to use prospection to form vivid mental representations of future outcomes^8^. Following earlier theoretical work^9^ initial empirical work confirmed that engagement in episodic future thinking can reduce temporal discounting behavior^10^. This effect has since then been replicated numerous times using a range of different tasks and experimental manipulations, as outlined in a recent meta-analysis^11^.

Episodic future thinking has been shown to affect temporal discounting in a variety of experimental designs^7,11^. Previous work from our group has focused on trial-wise^10,12,13^ and block-wise^14,15^ presentation of episodic cues during temporal discounting. In this experimental design, control trials involve choices between smaller-but-sooner and larger-but-later rewards. In some trials (episodic trials), the larger-but-later reward is additionally enriched by verbal episodic cues (tags) that serve as reminders of subject-specific events scheduled for the respective future time point associated with the delayed reward. In our trial-wise design, episodic and control trials are randomly intermixed^10,12,13^, whereas in the block.-wise design, blocks of episodic and control trials are completed separately in the same experimental session^14,15^. We investigated this effect in a number of previous studies summarized in Table 1 with n=204 participants in total. However, because modeling methods have continued to evolve, in our previous studies we have applied a range of different analytical approaches and model estimation schemes. In Peters & Buchel (2010), we fit hyperbolic discounting functions to estimated indifference points, separately for each participant and experimental condition, an approach that can be associated with some methodological problems^16^, e.g. this approach confounds goodness-of-fit (R^2^) with the discount rate^16^. In Bromberg et al. (2017) and Sasse et al. (2015, 2017), we applied Maximum Likelihood estimation using softmax action selection^17^, and fitted models separately to the data from each subject and condition. In Wiehler et al. (2017), we used hierarchical Bayesian parameter estimation, and assumed separate group-level distributions per group (gamblers vs. controls) and condition (episodic vs. control). It is clear that effect size estimates obtained from these different analytical approaches are not readily comparable, which poses a problem for e.g. meta-analyses of factors influencing discounting behavior^11^ or future power analyses that depend on the availability of comparable effect size estimates for different subject populations or age groups.

We therefore re-analyzed all our previously published data sets using the identical Bayesian estimation framework and using exactly the same hierarchical Bayesian model, yielding effect size estimates that are unconfounded by differences in the applied analytical approaches.

## Methods

We re-analyzed six data sets from five previously published studies on the effects of trial-wise and block-wise episodic cues on temporal discounting behavior (see Table 1).

**Table 1.**
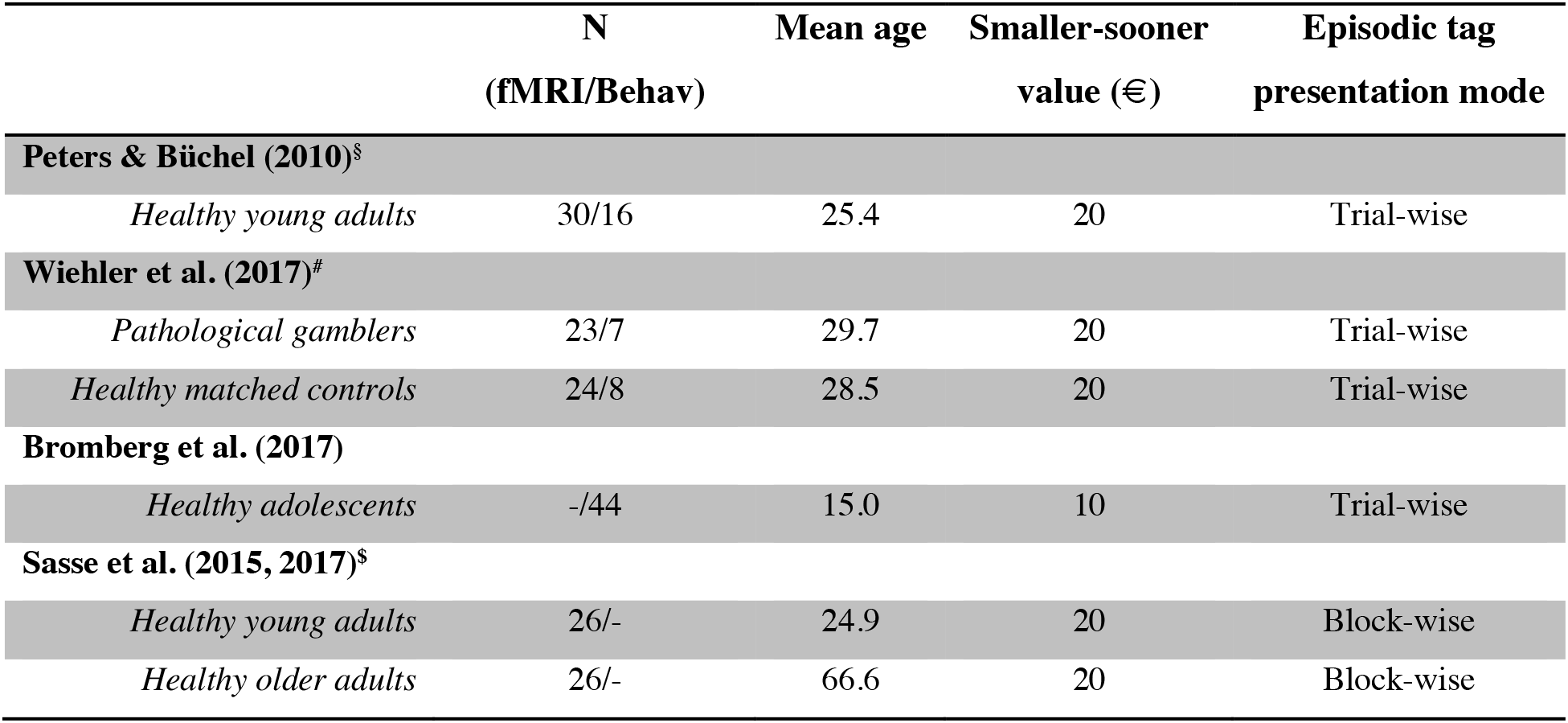
Overview of the included data sets and temporal discounting task details. ^§^Data from Experiment 1 (n=30) and the Experiment 2 (n=16) were pooled. ^#^Data from n=23 (n=24) gamblers (controls) that underwent fMRI was pooled with data from n=7 (n=8) gamblers (controls) that only performed a behavioral version of the task. ^$^In addition to the participants reported in Sasse et al. (2015, 2017), data from participants that were excluded from the fMRI analyses due to excessive motion were included in the present re-analysis.

In the control condition, participants made repeated choices between a smaller-but-sooner (SS) reward available immediately, and larger-but-later (LL) rewards.

In the trial-wise presentation studies^10,12,13^, in episodic trials subjects were additionally presented with subject-specific episodic event cues (episodic tags) referring to events planned at the respective time of delivery of the LL reward. These events were obtained for each individual participant in pre-experimental interviews and always referred to real and subject-specific events. Experiment 2 from Peters & Buchel (2010) additionally included an “unspecific” condition with hypothetical event cues. Due to lack of comparability with the other data sets, these data were not included here. In the trial-wise studies, participants completed 112 trials for each condition, and episodic and control trials were randomly intermixed.

In the block-wise presentation studies^14,15^, control blocks (two blocks of thirty six trials each) consisted of trials without episodic information. Episodic familiar blocks (two blocks of thirty-six trials each) involved the additional reference to a personally familiar event (individualized, each block with one distinct cue, e.g. “meeting with mum”). Data from the episodic unfamiliar condition^14,15^ included a cue referring to an unfamiliar episode (e.g. “meeting chancellor Merkel”). These data were not included here for lack of comparability with the trial-wise experimental designs outline above, which always included personally familiar episodes.

For further details regarding trial construction and timing we refer the reader to the original publications (see Table 1).

## Computational modeling

### Temporal discounting model

We applied a simple single-parameter hyperbolic discounting model to account for how value changes as a function of delay:

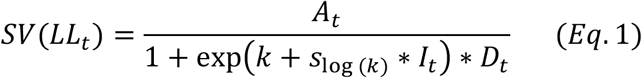

Here, *A_t_* is the numerical reward amount of the LL option on trial *t, D_t_* is the LL delay in days on trial *t* and *I_t_* is an indicator variable that takes on a value of 1 for trials from the episodic condition (including episodic tags) and 0 for trials from the control condition (without episodic tags). The value function has two free parameters: *k* is the hyperbolic discounting rate from the control condition (modeled in log-space) and *s_k_* is a coefficient modeling the degree of change in discounting for episodic vs. control trials.

We then used softmax action selection to model choice probabilities as a sigmoid function of value differences^18^:

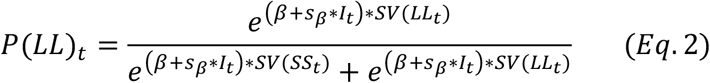

Here, *SV* is the subjective value of the delayed reward according to Eq. 1 and *β* is an inverse temperature parameter, modeling choice stochasticity (for *β* = 0, choices are random and as *β* increases, choices become more dependent on the option values). The value of the immediate smaller-sooner reward *SV*(*SS_t_*) was fixed throughout the experiments, but differed between studies (see Table 1). *I_t_* is again a dummy predictor coding for the episodic condition, and *s_β_* models the effect of the episodic condition on *β.*

### Data sets and participants

Model fitting was performed separately for each of the six datasets (see Table 1). For the Peters & Buchel (2010) and Wiehler et al. (2017) studies, we pooled data across participants that underwent fMRI, and participants that performed the same task without imaging (i.e. Experiment 1 and the temporally specific condition of Experiment 2 for Peters & Buchel (2010); behavioral pilot subjects and fMRI participants for Wiehler et al. (2017)). Note that we excluded one participant from the gambling group of the Wiehler et al. (2017) study, who made only a very small number of larger-later choices. Inclusion of this participant resulted in the hierarchical model that included within-subject changes in the softmax *β* parameter (Eq. 2) to fail to converge. For the Sasse et al. (2015, 2017) data, we additionally included the data from participants that were excluded from the original analyses due to excessive motion during fMRI (n=3 for Sasse et al., 2015; n=4 for Sasse et al., 2017), which is the reason for the discrepancy in sample sizes between Table 1 and the original papers.

**Table 2.** Overview of priors for group means.

| Parameter | Group-level prior ($\mu$) |
| --- | --- |
| $\log(k)$ | Uniform (-20, 3) |
| $s_{\log(k)}$ | Gaussian (0, 2) |
| $\beta$ | Uniform (0,10) |
| $s_{\beta}$ | Gaussian (0, 2) |

### Hierarchical Bayesian models

We analyzed each data set of Table 1 separately. Models were fit to all trials from all participants in a hierarchical Bayesian framework using Markov Chain Monte Carlo as implemented in JAGS 4.2.0^19^ using the *matjags* interface for Matlab © (The Mathworks). We applied the same hierarchical model for each data set. Parameter values for each participant were drawn from group-level Gaussian distributions, the mean and precision of which were estimated from the data. For group-level precision parameters, we used Gamma distributed priors. For group-level means of *log*(*k*) and *β,* we used uniform priors defined over numerically plausible parameter ranges (see Table 2). For group-level means of effects of the episodic condition (*s_log(k)_*, *s_β_*) we used Gaussian priors centered at zero (see Table 2). JAGS model code is available on the Open Science Framework (https://osf.io/bkgfd/).

We ran two chains with a burn-in period of 150k samples and thinning factor of 2. 5k additional samples where then retained for further analysis. Chain convergence was confirmed using the Gelman-Rubinstein convergence diagnostic 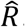, where we considered values of 1 ≤ 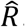 ≤ 1.01 as acceptable for all group-level and subject-level parameters. Evidence for an effect of the episodic future thinking manipulation was examined by computing Bayes Factors testing for directional effects^20,21^ on the posterior distributions of the group-level means of *s_log(k)_* and *s_β_*. We also report standardized effect sizes for all condition effects (Cohen’s *d*) which were calculated based on the means of the posterior means and precisions of *s_log(k)_* and *s_β_* (see Equations 1 and 2).

## Results

Mean changes for log(k) and *β* (*S_log(k)_, s_β_*) in the episodic condition are summarized in Table 3. Figure 1 shows posterior distributions for log(k) (upper panels) as well as changes in log(k) due to episodic cueing (lower panels) for all data sets.

**Table 3.** Overview of mean condition effects (change in model parameters from control to episodic condition) and Bayes Factors for directional effects (dBF: parameter reduction vs. increase). Standardized effect sizes (Cohen’s *d*) were calculated based on the estimated group-level posterior mean and precision parameters for *s*_log(*k*)_ (see Eq. 1) and s_β_ (see Eq. 2) from the hierarchical model. ^§^Data from Experiments 1 and the temporally specific condition of experiment 2 were pooled. ^#^Only data from the personally familiar episodic and control conditions were included (see Sasse et al., 2015, 2017).

| | Log (k) | | | Softmax $\beta$ | | |
| --- | --- | --- | --- | --- | --- | --- |
| | M (SD) | $d$ | dBF | M (SD) | $d$ | dBF |
| **Peters & Büchel (2010)<sup>§</sup>** |  |  |  |  |  |  |
| *Healthy young adults (n=46)* | -.108 (.207) | -.519 | 97.15 | -.004 (.059) | -.071 | 1.24 |
| **Wiehler et al. (2017)** |  |  |  |  |  |  |
| *Pathological gamblers (n=30)* | -.120 (.217) | -.552 | 34.69 | -.034 (.149) | -.223 | 4.43 |
| *Healthy controls (n=32)* | -.069 (.240) | -.287 | 6.52 | .020 (.071) | .284 | 0.26 |
| **Bromberg et al. (2017)** |  |  |  |  |  |  |
| *Adolescents (n=44)* | -.221 (.277) | -.799 | 2441.2 | .003 (.061) | .041 | 1.02 |
| **Sasse et al. (2015, 2017)<sup>#</sup>** |  |  |  |  |  |  |
| *Healthy young adults (n=26)* | -.280 (.578) | -.485 | 38.89 | .075 (.264) | .284 | 3.62 |
| *Healthy older adults (n=26)* | .006 (.315) | .018 | .91 | -.054 (.146) | -.367 | .115 |

**Figure 1.**
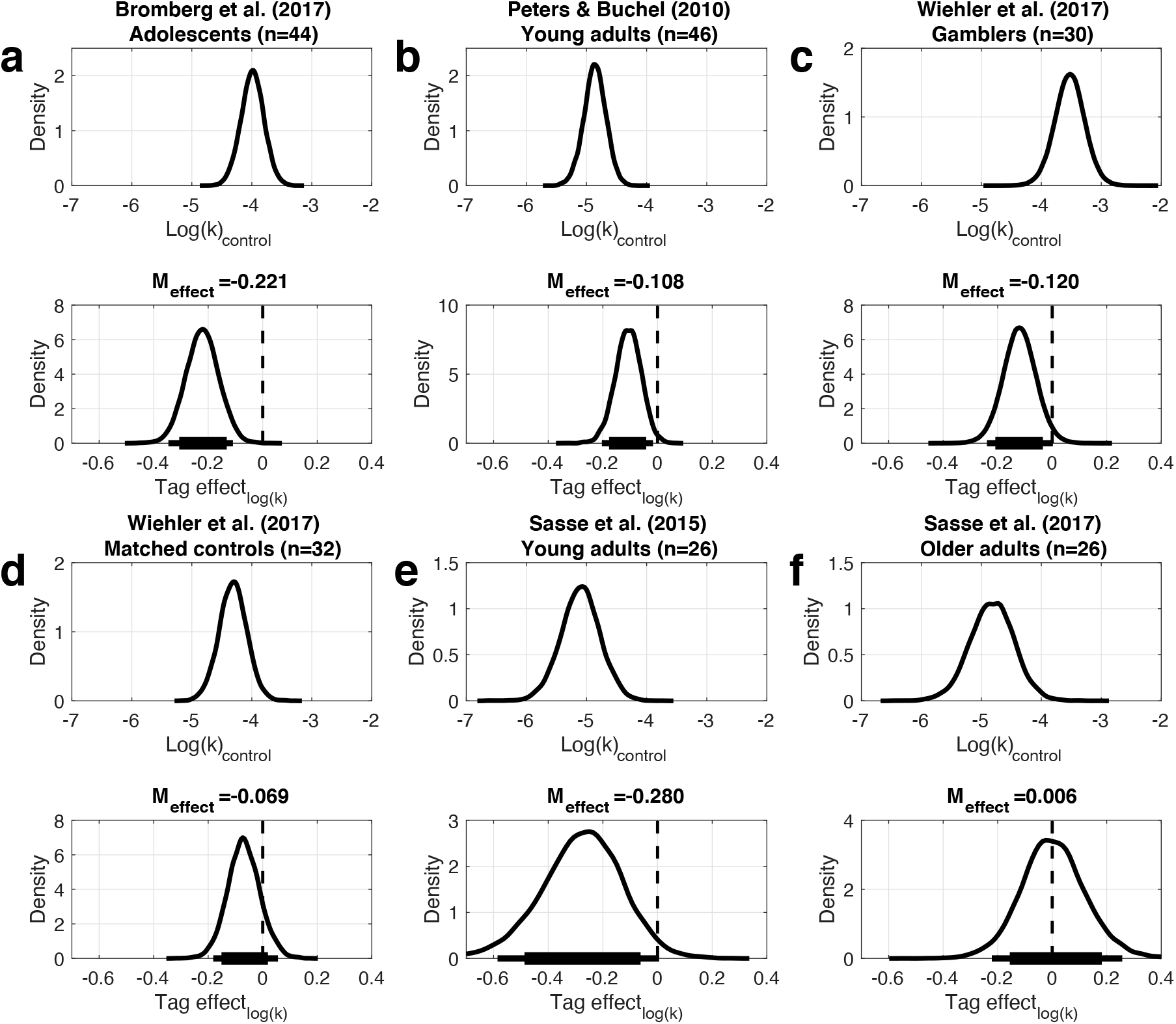
Posterior distributions of the hyperbolic discount rate log(k) in the control condition (upper panels) and the change in log(k) in the episodic vs. the control condition (*s*_log(*k*)_, lower panels). a) Healthy adolescents from Bromberg et al. (2017). b) Healthy young adults from Peters & Buchel (2010). c) Gambling disorder participants from Wiehler et al. (2017). d) Healthy matched controls from Wiehler et al. (2017). e) Healthy young adults from Sasse et al. (2015). f) Healthy older adults from Sasse et al. (2017). The solid (thin) horizontal lines denote 85% (95%) highest density intervals.

Across studies, the change in log(k) was consistently negative, with the older adults from Sasse et al. 2017 being the only exception with a posterior distribution that was centered at zero. However, effect sizes (mean changes) in log-space ranged from −.07 to −.280 (*d* ranged from .018 to −.799, Table 3 and Figure 1). There was also heterogeneity in this effect across participants. We illustrate this variability in Figure 2, where we plot posterior distributions of *s*_log(*k*)_ for each individual participant, separately for the six data sets.

In contrast, the observed mean changes in *β* (see Figure 3) were generally small and with inconsistent directionality across groups (see Table 3).

**Figure 2.**
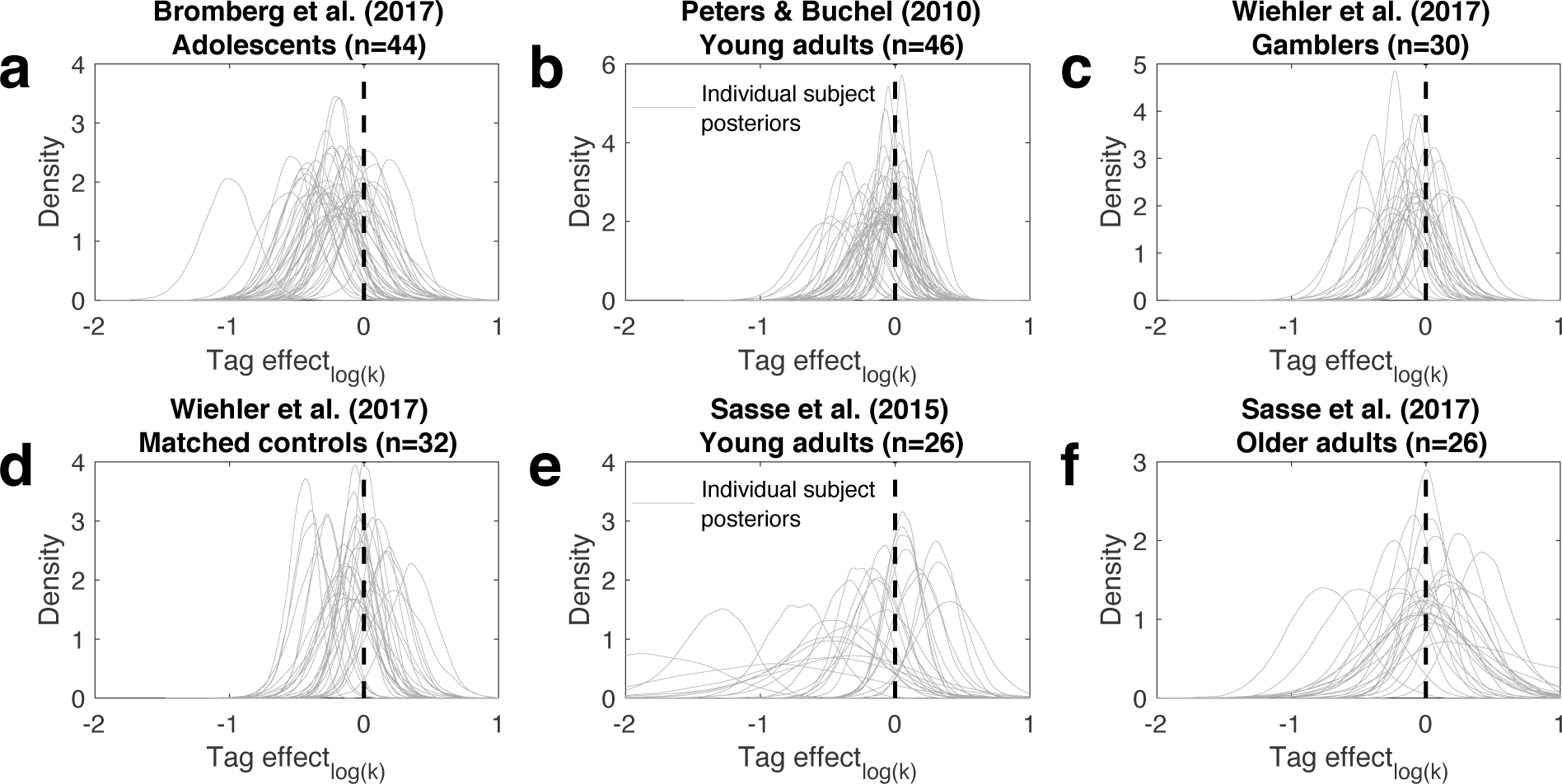
Individual subject posterior distributions of the change in log(k) in the episodic vs. the control condition (*s*_log (*k*)_). a) Healthy adolescents from Bromberg et al. (2017). b) Healthy young adults from Peters & Buchel (2010). c) Gambling disorder participants from Wiehler et al. (2017). d) Healthy matched controls from Wiehler et al. (2017). e) Healthy young adults from Sasse et al. (2015). f) Healthy older adults from Sasse et al. (2017).

**Figure 3.**
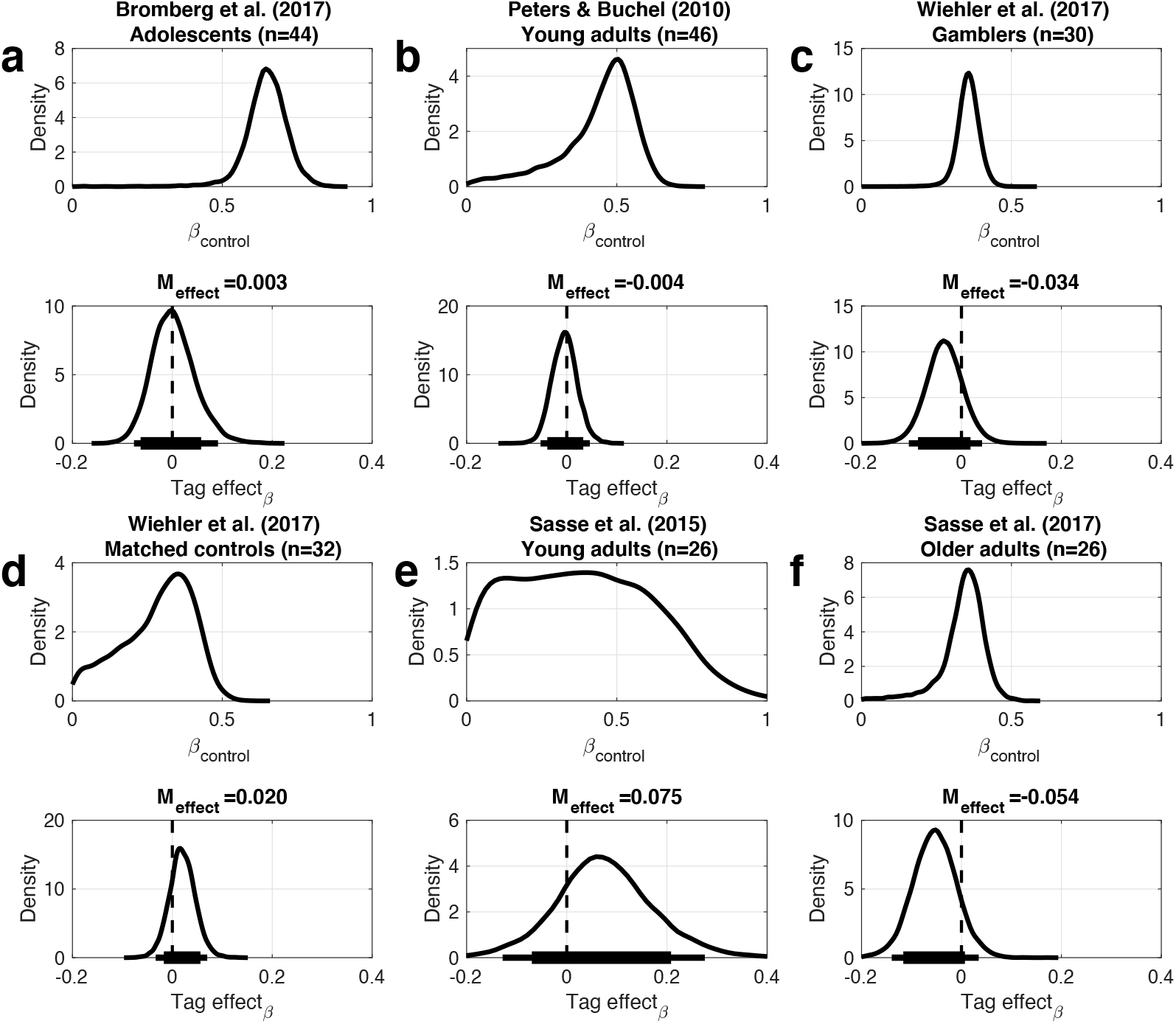
Posterior distributions of softmax inverse temperature parameter *β* in the control condition (top row) and the change in *β* in the episodic vs. the control condition (*s_β_*, bottom row). a) Healthy adolescents from Bromberg et al. (2017). b) Healthy young adults from Peters & Buchel (2010). c) Gambling disorder participants from Wiehler et al. (2017). d) Matched controls from Wiehler et al. (2017). e) Young adults from Sasse et al. (2015). f) Older adults from Sasse et al. (2017). The solid (thin) horizontal lines denote 85% (95%) highest density intervals.

## Discussion

Here we re-analyzed previously published data on episodic future thinking effects on temporal discounting (six data sets from five papers, n=204 subjects in total). Our results provide comparable effect size estimates across studies, and confirm robust and consistent effects of episodic future thinking on temporal discounting in designs involving trial-wise and block-wise presentations of episodic cues (tag effect) in most data sets.

Our analyses advance over previous analytical approaches in three ways. First, we accounted for the within-subject nature of the experimental design at the model estimation stage. That is, model parameters in the control condition were modeled as the baseline, and changes from that baseline due to episodic cueing were modeled as additive within-subject changes. There is generally a high level of both short-term and long-term stability in discount rates^22–25^. Modeling condition effects as within-subject changes from a baseline condition reduces the variability in the posterior distribution of the treatment effect (e.g.. *s*_log(*k*)_), compared to a model that estimates independent parameters or posterior distributions per condition, as we have done previously^10,12–15,10,12,13^. Second, our use of hierarchical Bayesian parameter estimation entails additional advantages. In hierarchical Bayesian estimation, individual participant’s parameters are assumed to be drawn from group-level Gaussian distributions, such that each participant’s parameters are informed and constrained by the distribution of parameters in the entire sample. This “partial pooling” or “shrinkage” can increase the robustness of the resulting estimates^26^. Finally, we have applied the exact same hierarchical model and estimation procedure across all six datasets. Consequently, the effect size estimates reported here are unconfounded by differences in model structure, priors, and/or estimation procedures, and therefore constitute the best available estimates of effect sizes for these experimental designs.

In contrast to some of our earlier work^10,13^, we additionally examined the degree to which episodic tags affected overall decision noise (softmax *β*). Note that potential changes in *β* could reflect differences in the best-fitting discounting model^17^ and/or unmodeled systematic influences on choice patterns, as well as the level unsystematic noise in the behavioral data. In contrast to episodic effects on log(k), which showed consistent directionality across studies and generally medium effect sizes, episodic cueing effects on *β* where generally smaller and showed inconsistent directionality across studies. Under some conditions, changes in decision noise can seemingly give rise to changes in discounting behavior^27–29^, an effect that depends on the individual level of discounting in relation to the space of choice options examined in a given experimental task. The fact that episodic thinking effects on *β* where generally small and of inconsistent directionality argues against an unspecific effect of episodic cues on overall choice patterns.

Our analysis revealed robust effects of episodic future thinking on temporal discounting in pathological gamblers. In our previous report^12^, we did not account for the within-subject nature of the design at the model estimation stage, which likely increased the variance in the observed group-level parameters, precluding us from accurately estimating the magnitude and variance of the episodic tag effect. Here we show that the effect size of the tag effect on log(k) in pathological gamblers is in fact of comparable magnitude to that observed in our previous study in healthy young participants^10^, while the effect is somewhat less pronounced in the healthy matched control group of the Wiehler et al. (2017) study. This means that, if anything, we have previously underestimated the magnitude of this effect in pathological gamblers. In the light of the fact that increases in temporal discounting are implicated in a range of psychiatric disorders^3,5,30^, this is a promising first finding. However, a central question that remains to be addressed by the field is whether experimental modulations of discounting behavior can yield clinically relevant behavioral changes^30^. In contrast to training-based interventions^31^, the present experimental design is likely not suited to induce long lasting changes in behavior. Nonetheless, our data show that in principle, future thinking can reduce temporal discounting in pathological gamblers, a clinical group characterized by high levels of impulsivity^4^. Furthermore, the observed changes in this clinical sample were similar in magnitude to those observed in healthy young adults. Future studies will likely build upon recent work that aimed to extend future thinking interventions to everyday decision-making^32–36^.

In contrast to the findings in healthy young adolescents and adults as well as gamblers, older adults showed no effect of future thinking on temporal discounting^15^. This was also shown in a recent paper from another group^37^. As discussed in detail in our previous paper^15^, in older adults effects of future thinking might depend on cognitive control abilities. We have shown that older adults with high levels of cognitive control still benefitted from future thinking, whereas this was not the case for older adults with relatively lower control abilities^15^. It remains to be seen whether similar moderation effects play a role in other age groups or populations.

This re-analysis of previously published data has a number of limitations. First, the experimental designs that we applied involved a separation of decision and response phases, as appropriate for fMRI studies. This precluded us from applying modeling approaches that leverage information contained in the response time (RT) distributions, as in some of our more recent work^38,39^. Second, comparison of the effect size estimates between the Bromberg et al. (2017) data set and the other studies is confounded by the fact that the smaller-sooner reference reward in that study consisted of 10€, whereas it was 20€ in the other data sets. Steeper discounting and/or a more pronounced effect of the episodic condition in adolescents could thus be partially attributable to a magnitude effect^40–43^.

Taken together, our re-analysis of six previously published data sets that examined the effects of episodic future thinking on temporal discounting provides comparable effect size estimates across studies. We hope this resource to be helpful for both future power analyses and for meta-analyses on contextual modulations of temporal discounting more generally.

## Acknowledgements

This work was funded by the DFG (grants PE1627/3-1 and PE1627/5-1 to J.P. and BR2877/2-2 to S.B.).

## References

1. Green, L. & Myerson, J. A discounting framework for choice with delayed and probabilistic rewards. Psychol Bull 130, 769–92 (2004).

2. Peters, J. & Büchel, C. The neural mechanisms of inter-temporal decision-making: understanding variability. Trends Cogn. Sci. 15, 227–239 (2011).

3. Bickel, W. K., Jarmolowicz, D. P., Mueller, E. T., Koffarnus, M. N. & Gatchalian, K. M. Excessive discounting of delayed reinforcers as a trans-disease process contributing to addiction and other disease-related vulnerabilities: Emerging evidence. Pharmacol Ther 134, 287–97 (2012).

4. Wiehler, A. & Peters, J. Reward-based decision making in pathological gambling: the roles of risk and delay. Neurosci. Res. 90, 3–14 (2015).

5. Amlung, M. et al. Delay Discounting as a Transdiagnostic Process in Psychiatric Disorders: A Meta-analysis. JAMA Psychiatry (2019) doi:10.1001/jamapsychiatry.2019.2102.

6. Koffarnus, M. N., Jarmolowicz, D. P., Mueller, E. T. & Bickel, W. K. Changing delay discounting in the light of the competing neurobehavioral decision systems theory: a review. J. Exp. Anal. Behav. 99, 32–57 (2013).

7. Lempert, K. M. & Phelps, E. A. The Malleability of Intertemporal Choice. Trends Cogn. Sci. 20, 64–74 (2016).

8. Schacter, D. L., Benoit, R. G. & Szpunar, K. K. Episodic Future Thinking: Mechanisms and Functions. Curr. Opin. Behav. Sci. 17, 41–50 (2017).

9. Boyer, P. Evolutionary economics of mental time travel? Trends Cogn Sci 12, 219–24 (2008).

10. Peters, J. & Büchel, C. Episodic Future Thinking Reduces Reward Delay Discounting through an Enhancement of Prefrontal-Mediotemporal Interactions. Neuron 66, 138–148 (2010).

11. Rung, J. M. & Madden, G. J. Experimental reductions of delay discounting and impulsive choice: A systematic review and meta-analysis. J. Exp. Psychol. Gen. 147, 1349–1381 (2018).

12. Wiehler, A., Petzschner, F. H., Stephan, K. E. & Peters, J. Episodic Tags Enhance Striatal Valuation Signals during Temporal Discounting in pathological Gamblers. eNeuro 4, (2017).

13. Bromberg, U., Lobatcheva, M. & Peters, J. Episodic future thinking reduces temporal discounting in healthy adolescents. PloS One 12, e0188079 (2017).

14. Sasse, L. K., Peters, J., Büchel, C. & Brassen, S. Effects of prospective thinking on intertemporal choice: The role of familiarity. Hum. Brain Mapp. 36, 4210–4221 (2015).

15. Sasse, L. K., Peters, J. & Brassen, S. Cognitive Control Modulates Effects of Episodic Simulation on Delay Discounting in Aging. Front. Aging Neurosci. 9, 58 (2017).

16. Johnson, M. W. & Bickel, W. K. An algorithm for identifying nonsystematic delay-discounting data. Exp Clin Psychopharmacol 16, 264–74 (2008).

17. Peters, J., Miedl, S. F. & Büchel, C. Formal Comparison of Dual-Parameter Temporal Discounting Models in Controls and Pathological Gamblers. PLoS ONE 7, e47225 (2012).

18. Sutton, R. S. & Barto, A. G. Reinforcement Learning: An Introduction. (MIT Press, 1998).

19. Plummer, M. JAGS: A program for analysis of Bayesian graphical models using Gibbs sampling. in Proceedings of the 3rd international workshop on distributed statistical computing vol. 124 125 (Technische Universit at Wien, 2003).

20. Marsman, M. & Wagenmakers, E.-J. Three Insights from a Bayesian Interpretation of the One-Sided P Value. Educ. Psychol. Meas. 77, 529–539 (2017).

21. Pedersen, M. L., Frank, M. J. & Biele, G. The drift diffusion model as the choice rule in reinforcement learning. Psychon. Bull. Rev. 24, 1234–1251 (2017).

22. Enkavi, A. Z. et al. Large-scale analysis of test-retest reliabilities of self-regulation measures. Proc. Natl. Acad. Sci. U. S. A. 116, 5472–5477 (2019).

23. Kirby, K. N. One-year temporal stability of delay-discount rates. Psychon Bull Rev 16, 457–62 (2009).

24. Ohmura, Y., Takahashi, T., Kitamura, N. & Wehr, P. Three-month stability of delay and probability discounting measures. Exp Clin Psychopharmacol 14, 318–28 (2006).

25. Peters, J. & Buchel, C. Overlapping and Distinct Neural Systems Code for Subjective Value during Intertemporal and Risky Decision Making. J. Neurosci. 29, 15727–15734 (2009).

26. Farrell, S. & Lewandowsky, S. Computational modeling of cognition and behavior. (Cambridge University Press, 2018).

27. Hinson, J. M., Jameson, T. L. & Whitney, P. Impulsive decision making and working memory. J Exp Psychol Learn Mem Cogn 29, 298–306 (2003).

28. Hinson, J. M. & Whitney, P. Working memory load and decision making: A reply to Franco-Watkins, Pashler, and Rickard (2006). J. Exp. Psychol. Learn. Mem. Cogn. 32, 448–450 (2006).

29. Franco-Watkins, A. M., Pashler, H. & Rickard, T. C. Does working memory load lead to greater impulsivity? Commentary on Hinson, Jameson, and Whitney (2003). J. Exp. Psychol. Learn. Mem. Cogn. 32, 443–447; discussion 448-450 (2006).

30. Lempert, K. M., Steinglass, J. E., Pinto, A., Kable, J. W. & Simpson, H. B. Can delay discounting deliver on the promise of RDoC? Psychol. Med. 49, 190–199 (2019).

31. Bickel, W. K., Yi, R., Landes, R. D., Hill, P. F. & Baxter, C. Remember the Future: Working Memory Training Decreases Delay Discounting Among Stimulant Addicts. Biol. Psychiatry 69, 260–265 (2011).

32. Daniel, T. O., Said, M., Stanton, C. M. & Epstein, L. H. Episodic future thinking reduces delay discounting and energy intake in children. Eat. Behav. 18, 20–24 (2015).

33. Daniel, T. O., Stanton, C. M. & Epstein, L. H. The future is now: reducing impulsivity and energy intake using episodic future thinking. Psychol. Sci. 24, 2339–2342 (2013).

34. O’Neill, J., Daniel, T. O. & Epstein, L. H. Episodic future thinking reduces eating in a food court. Eat. Behav. 20, 9–13 (2016).

35. Stein, J. S. et al. Unstuck in time: episodic future thinking reduces delay discounting and cigarette smoking. Psychopharmacology (Berl.) 233, 3771–3778 (2016).

36. Daniel, T. O., Stanton, C. M. & Epstein, L. H. The future is now: Comparing the effect of episodic future thinking on impulsivity in lean and obese individuals. Appetite 71, 120–125 (2013).

37. Mok, J. N. Y. et al. Is it time? Episodic imagining and the discounting of delayed and probabilistic rewards in young and older adults. Cognition 199, 104222 (2020).

38. Peters, J. & D’Esposito, M. The drift diffusion model as the choice rule in intertemporal and risky choice: a case study in medial orbitofrontal cortex lesion patients and controls. bioRxiv 642587 (2019) doi:10.1101/642587.

39. Wagner, B., Clos, M., Sommer, T. & Peters, J. Dopaminergic modulation of human inter-temporal choice: a diffusion model analysis using the D2-receptor-antagonist haloperidol. bioRxiv 2020.02.13.942383 (2020) doi:10.1101/2020.02.13.942383.

40. Ballard, I. C. et al. More Is Meaningful: The Magnitude Effect in Intertemporal Choice Depends on Self-Control. Psychol. Sci. 28, 1443–1454 (2017).

41. Green, L., Myerson, J., Oliveira, L. & Chang, S. E. Delay discounting of monetary rewards over a wide range of amounts. J Exp Anal Behav (2013).

42. Green, L., Myerson, J. & McFadden, E. Rate of temporal discounting decreases with amount of reward. Mem. Cognit. 25, 715–723 (1997).

43. Mellis, A. M., Woodford, A. E., Stein, J. S. & Bickel, W. K. A second type of magnitude effect: Reinforcer magnitude differentiates delay discounting between substance users and controls. J. Exp. Anal. Behav. 107, 151–160 (2017).

